# Improving ATAC-seq Data Analysis with AIAP, a Quality Control and Integrative Analysis Package

**DOI:** 10.1101/686808

**Authors:** Shaopeng Liu, Daofeng Li, Cheng Lyu, Paul Gontarz, Benpeng Miao, Pamela Madden, Ting Wang, Bo Zhang

**Author notes:** The authors wish it to be known that, in their opinion, the first three authors should be regarded as joint First Authors.

## Abstract

ATAC-seq is a technique widely used to investigate genome-wide chromatin accessibility. The recently published Omni-ATAC-seq protocol substantially improves the signal/noise ratio and reduces the input cell number. High-quality data are critical to ensure accurate analysis. Several tools have been developed for assessing sequencing quality and insertion size distribution for ATAC-seq data; however, key quality control (QC) metrics have not yet been established to accurately determine the quality of ATAC-seq data. Here, we optimized the analysis strategy for ATAC-seq and defined a series of QC metrics, including reads under peak ratio (RUPr), background (BG), promoter enrichment (ProEn), subsampling enrichment (SubEn), and other measurements. We incorporated these QC tests into our recently developed ATAC-seq Integrative Analysis Package (AIAP) to provide a complete ATAC-seq analysis system, including quality assurance, improved peak calling, and downstream differential analysis. We demonstrated a significant improvement of sensitivity (20%~60%) in both peak calling and differential analysis by processing paired-end ATAC-seq datasets using AIAP. AIAP is compiled into Docker/Singularity, and with one command line execution, it generates a comprehensive QC report. We used ENCODE ATAC-seq data to benchmark and generate QC recommendations, and developed qATACViewer for the user-friendly interaction with the QC report.

## INTRODUCTION

To regulate the transcription of a eukaryotic genome, chromatin must remain in an accessible state to allow binding of transcription factors and initiation of transcription activation[1–4]. Several sequencing-based methods have been developed to assess chromatin accessibility and nucleosome positioning, including DNase-seq[5], FAIRE-seq[6], MNase-seq[7], and the recently developed ATAC-seq (assay for transposase-accessible chromatin followed by sequencing)[8]. ATAC-seq can detect the accessible regions of a genome by identifying open chromatin regions using a prokaryotic Tn5-transposase[8, 9], and the technology features an easy experimental protocol, a reduced requirement of input material, and a high signal/noise ratio. These unique advantages have propelled ATAC-seq technology to quickly become a widely used method to define chromatin accessibility, especially in several large consortiums focusing on functional genomics profiling, including ENCODE[10], TaRGET II[11], and IHEC[12].

The ATAC-seq analysis strategy is primarily adopted from ChIP-seq data analysis. After aligning sequencing reads to the genome, peak-calling tools, such as MACS2[13], are commonly used to identify highly enriched ATAC-seq signals across the genome. Unlike ChIP-seq, an ATAC-seq experiment does not normally require input control. Thus, accurately assessing the quality of ATAC-seq data is a critical step influencing downstream analysis. Several software packages were developed for ATAC-seq quality control (QC) and data analysis[14–16]. These tools provide general QC metrics of sequencing data, including read quality score, sequencing depth, duplication rate, and library insert fragment size distribution. Many tools also provide analysis functions, including foot-printing analysis, motif analysis, and library complexity analysis.

Here, we present AIAP (ATAC-seq Integrative Analysis Package), a software package containing an optimized ATAC-seq data QC and analysis pipeline. Along with general QC metrics, such as library insert fragment size distribution, we specifically introduced a series of QC metrics for ATAC-seq, including reads under peak ratio (RUPr), background (BG), promoter enrichment (ProEn), subsampling enrichment (SubEn), and other measurements. By applying AIAP, we demonstrated a significant improvement in both peak calling and differential analysis by processing the paired-end sequencing data in single-end mode: more than 20% of ATAC-seq peaks can be identified using AIAP, and over 30% more differentially accessible regions (DARs) can be identified by AIAP in downstream analysis. We applied AIAP to reanalyse 54 mouseENCODE[17] ATAC-seq datasets and determined the general QC recommendation for ATAC-seq data analysis. We also developed qATACViewer, a visualization tool included in AIAP, for user-friendly visualization of QC reports. AIAP is compiled into a Docker/Singularity image to allow maximized compatibility on different operating systems and computing platforms. The software, source code, and documentation are freely available at https://github.com/Zhang-lab/ATAC-seq_QC_analysis.

## METHODS

Here, we describe AIAP, **A**TAC-seq **I**ntegrative **A**nalysis **P**ackage, for processing and analysing ATAC-seq data. The AIAP workflow typically consists of four steps, as shown in **Figure 1**: (i) Data Processing; (ii) QC; (iii) Integrative Analysis; and (iv) Data Visualization. Below, we introduce the technical details of AIAP. Detailed documentation is available at https://github.com/Zhang-lab/ATAC-seq_QC_analysis/.

**Figure 1.**
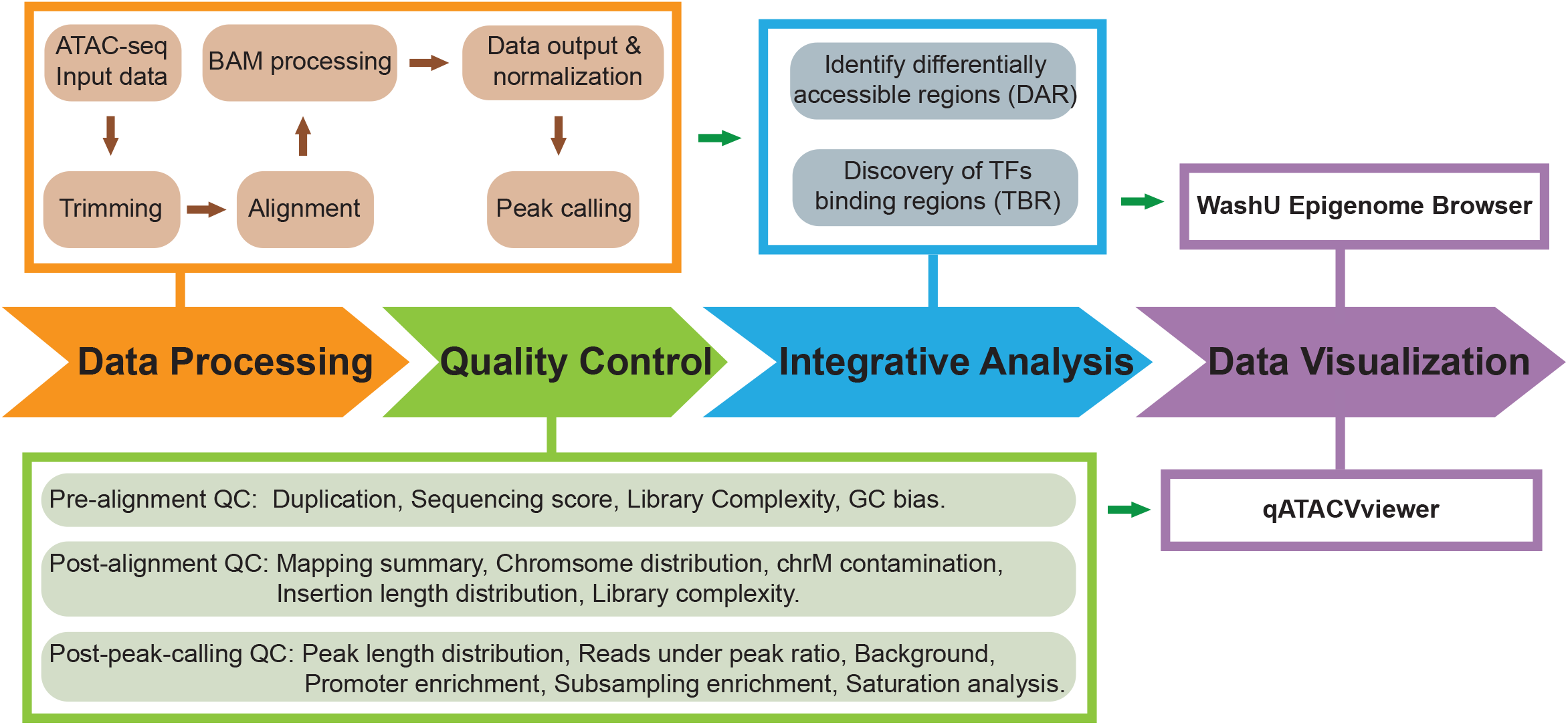
Schematic representation of ATAC-seq Integrative Analysis Package (AIAP). The schema reports the four analytical steps, namely, Data Processing, Quality Control, Integrative Analysis, and Data Visualization.

### Data processing

The data processing step first configures the working path. The ATAC-seq paired-end (PE) raw-read FASTQ files are trimmed by cutadapt and aligned to the reference genome by *bwa*[18]. The bam file is further processed by *methylQA*[19] in the ATAC mode. The methylQA first filters unmapped and low-quality mapped PE reads and then identifies the Tn5 insertion position at each read end by shifting +4 bp/−5 bp on the positive/negative strands. *methylQA* further extends 75 bp in both directions around the Tn5 insertion position to create two pseudo single-end (SE) mapped reads with 150 bp length in PE(asSE) mode. Next, AIAP compiles different files for downstream analysis (.bed files) and normalized visualization (.bigWig files). The bed file is used to perform peak calling by *MACS2*[13] with a q-value cutoff of 0.01 and the following setting: --*keep-dup 1000 --nomodel --shift 0 --extsize 150*.

### QC

AIAP performs a series of quality checking steps before and after alignment. AIAP calls *fastQ* to check the sequencing quality, duplication rate, and GC bias before alignment. After alignment, AIAP generates the mapping stat summary, chromosome distribution of uniquely mapped reads, chrM contamination rate, insertion fragment length distribution, and library complexity. AIAP also performs a series of post-peak-calling quality checks, including peak width distribution, read under peak ratio, background, and enrichment calculation (promoter and subsampling). AIAP also provides saturation analysis, promoter peak distribution, and signal ranking analysis. AIAP reports the quality metrics in a JSON file, which can be visualized using qATACViewer. We defined the key QC metrics as follows:

#### Alignment QC

*Non-redundant uniquely mapped reads*: the reads that are uniquely mapped to the reference genome after removing redundancy.

*Chromosome distribution/ChrM contamination rate*: distribution of uniquely mapped reads across all chromosomes. The number of uniquely mapped reads on ChrM is used as a QC metric to measure the quality of the ATAC-seq library.

*Library insert size distribution*: insertion size distribution is measured as the length of DNA fragment defined by non-redundant uniquely mapped read-pairs.

*Library complexity*: library complexity is estimated in both duplication rate and predicted yield of distinct reads generated by preSeq (https://github.com/smithlabcode/preseq).

*Useful single ends*: each end of a non-redundant uniquely mapped read pair will be shifted +4 bp/−5 bp on the positive/negative strands and then further extended 75 bp in both directions around the Tn5 insertion position.

#### Peak-calling QC

*Reads under the peak ratio* (*RUPr*): percentage of all useful ends (E) that fall into the called peak regions with at least 50% overlap. RUPr is calculated as follows:

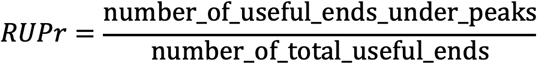

*Background*: Fifty thousand genomic regions (500bp each) are randomly selected from the genome outside of ATAC-seq peaks. The ATAC-seq signal in each region is calculated as reads per kilobase per million mapped reads (RPKM). The percentage of all such regions with the ATAC-seq signal over the theoretical threshold (RPKM=0.377) is considered high-background and used as a QC metric to indicate the background noise.

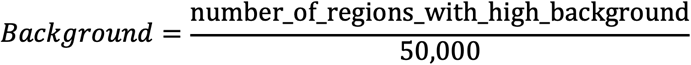

*Promoter enrichment* (*ProEn*): the promoters of active genes provide a positive control for open chromatin regions. The ATAC-seq useful ends (E) enriched on detected promoters (overlapping ATAC-seq peaks) are used as a QC metric to measure the signal enrichment calculated as follows:

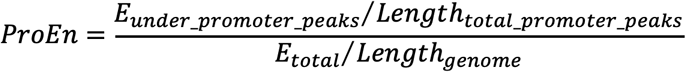

*Subsampling enrichment* (*SubEn*): the ATAC-seq signal (useful ends) enriched on the detected ATAC-seq peaks is used as a QC metric to measure the signal enrichment at the genome-wide level. To avoid sequencing-depth bias, 10 million useful ends (E) are sampled from the complete dataset, and peak calling is performed to identify the open chromatin regions. SubEn is calculated after 10 million pseudo counts are added into the calculation as background:

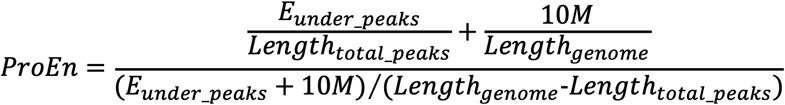

*Saturation Analysis*: MACS2 is used to call narrow peaks for a series of subsampling from complete useful ends (10% step). The length of identified peaks covering genomic regions at each subsampling are used to calculate the recovery (percentage) of complete peaks covering genomic regions when using complete useful ends.

*Signal ranking analysis*: the ATAC-seq peak signals are ranked, and the percentage of promoter peaks in each quantile is determined.

### Integrative analysis

AIAP includes two downstream analysis components: 1). Analysis of DARs between two conditions and 2). Discovery of transcription factor binding regions. AIAP calculates the read counts for all peaks identified under all conditions after peak calling, and a pair-wise comparison is performed by querying the R package DESeq2[20] based on the design table. AIAP will further identify potential TF binding regions under ATAC-seq peaks by implementing the Wellington algorithm[21].

### Data visualization

AIAP generates a collection of files for visualizing the ATAC-seq data on a genome browser[22–24], including the normalized signal density file (normalized to 10M total reads) in bigwig format, the Tn5 insertion positions file in bigwig format, the peak file in bed format, and the footprint positions file in bed format. AIAP generates a JSON QC report that can be visualized with the embedded qATACviewer [**Supplementary Figure 1**].

### DNase hypersensitive sites (DHSs), histone signal calculation

The raw data FASTQ files of ATAC-seq and histone ChIP-seq were downloaded from ENCODE data portal (https://www.encodeproject.org/), and listed in Supplementary Table 1. The ATAC-seq FASTQ files were processed by AIAP as described above. The ChIP-seq FASTQ files were aligned to the mouse genome (mm10 assembly) and were further processed by methylQA. The processed methylation calling of whole-genome bisulfite sequencing (WGBS) data was downloaded from the ENCODE data portal. The averaged signals of ATAC-seq, ChIP-seq, and WGBS were calculated at 100 bp windows within 5 kb around the centre of the open chromatin regions (OCRs) and were plotted in R.

### Differentially Accessible Regions (DARs) identification

The DARs of each tissue were identified between different mouse developmental stages (embryonic 11.5 day and postnatal 0 day) to evaluate the performance of AIAP. The OCRs generated by AIAP were used as test regions, the read counts were calculated in both PE(asSE) and PE(noShift) modes, and the DARs were identified as described above method with adjusted p-value cut-off less than 0.01 and absolute log2-transformed fold-change over 1.

## RESULTS

### Defining the QC metrics of ATAC-seq data

Conducting QC checks at different steps of data processing and correctly interpreting QC metrics are crucial to ensure a successful and meaningful analysis. Different QC metrics report important information regarding different aspects of genomic data; thus, it is essential to define the key QC metrics for ATAC-seq data before performing an analysis. In addition to the traditional QC metrics shown in **Figure 1**, we specifically chose reads under peak ratio (RUPr), background, and promoter enrichment (ProEn) as key QC metrics to measure the quality of ATAC-seq data. RUPr is an essential QC metric for ChIP-seq experiments[25] and is widely adopted to measure ATAC-seq data. The ENCODE consortium recommends that at least 20% of non-redundant uniquely mapped reads be located in peak regions. A higher RUPr usually indicates a high signal-to-noise ratio. Similar to RUPr, a higher promoter enrichment (ProEn) also indicates a high signal-to-noise ratio. ProEn is calculated to indicate the enrichment of the ATAC-seq signal over gene promoters, which are usually in open chromatin across different tissue and cell types. We used the ENCODE ATAC-seq data as a benchmark, and we determined that RUPr and ProEn directly reflect the quality of ATAC-seq at comparable sequencing depths [**Figure 2A**]. The ATAC-seq peaks are sharper and stronger when the ATAC-seq data have high RUPr and ProEn values, suggesting better signal enrichment and better quality in the ATAC-seq experiment.

**Figure 2.**
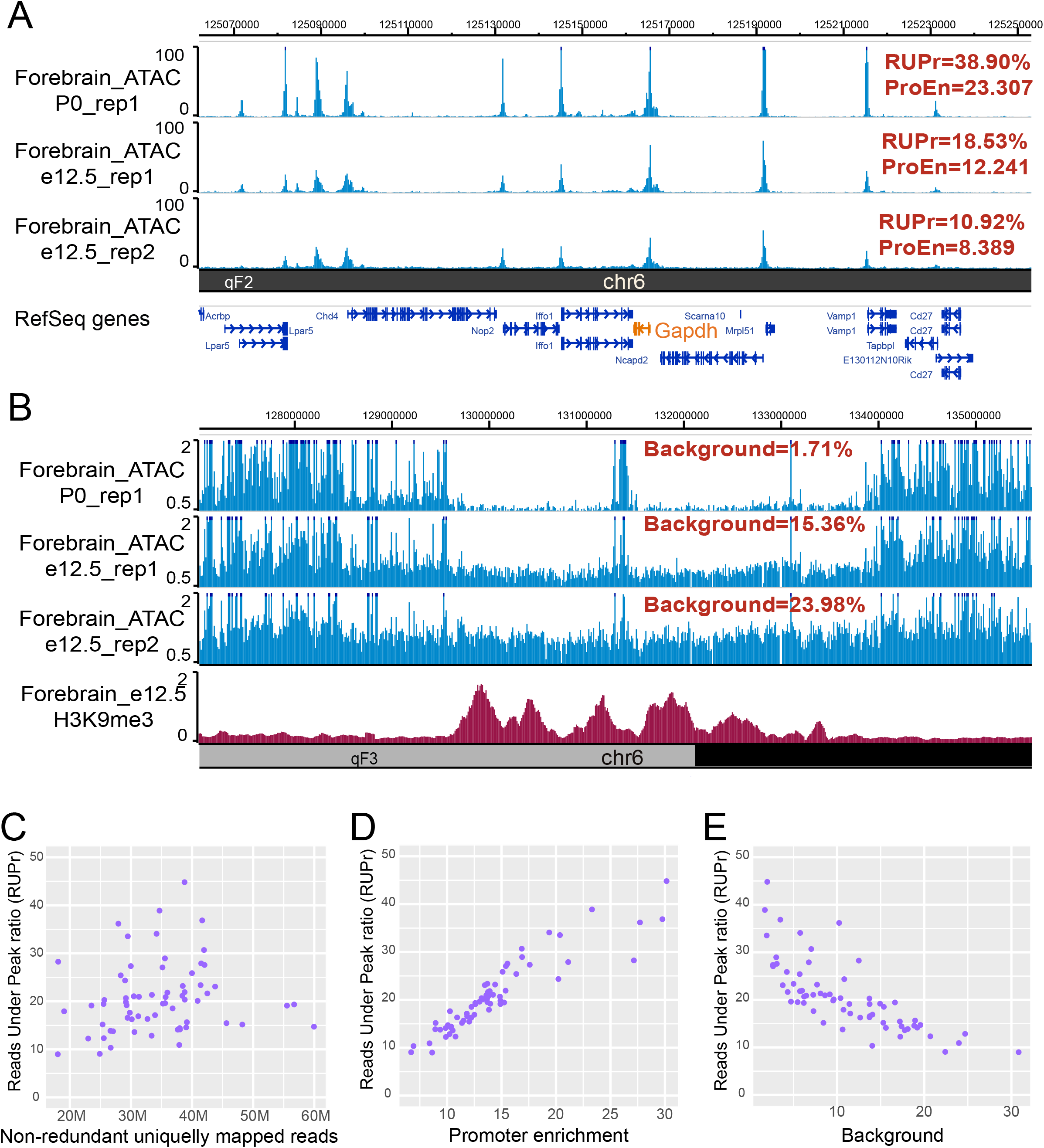
Key QC metrics of ATAC-seq. **A).** Three normalized (to 10M) ATAC-seq datasets with distinct RUPr and ProEn were visualized on the WashU Epigenome Browser. **B).** Three normalized (to 10M) ATAC-seq datasets (same as A) with distinct background values were visualized on the WashU Epigenome Browser. The centre regions were highly enriched for repressive H3K9me3 histone modification in the mouse forebrain. **C).** Scatter-plot between non-redundant uniquely mapped reads and read under peak ratio (RUPr). **D).** Scatter-plot between promoter enrichment (ProEn) and RUPr. **E).** Scatter-plot between background and RUPr.

We further defined “background” to directly measure the background noise level in the ATAC-seq experiment. We randomly selected 50,000 genomic regions (size: 500 bp) from regions of the genome that did not overlap with ATAC-seq peaks after peak calling. The ATAC-seq signals over each region were calculated as RPKM, and the regions with an ATAC-seq signal over a theoretical threshold (RPKM=0.377) were considered high-background regions. The percentage of high-background regions within 50,000 randomly selected genomic regions was used as a QC metric to measure the background noise of the ATAC-seq data. We noticed that different background noise levels were directly reflected by the QC metric of background, especially in the heterochromatin regions, which are enriched with H3K9me3 signals [**Figure 2B**].

To further explore the QC metrics, we used AIAP to process 54 ATAC-seq datasets generated by the ENCODE consortium [**Supplementary Table 2**]. We specifically checked the key metrics, including RUPr, ProEn, and background. We noticed that these metrics were not dependent on sequencing depth [**Figure 2C**, **Supplementary Figure 2**]. The RUPr was positively correlated with ProEn, proving an accurate measurement of signal enrichment in the ATAC-seq data [**Figure 2D**]. The background was negatively correlated with RUPr, providing a measurement of the noise level in the ATAC-seq data [**Figure 2E**].

### AIAP improved the sensitivity of discovering OCRs

To define OCRs, peak-calling strategies adopted from ChIP-seq analysis are widely used to analyse ATAC-seq data. However, unlike ChIP-seq data, an ATAC-seq experiment does not have input control and is usually sequenced with the PE sequencing method to profile the size of DNA fragments. The uniquely aligned PE reads have been used to call open chromatin peaks after alignment in many studies[26–32]. After detecting the distribution of reads with different lengths under the peak regions, we noticed that the medium fragments and long fragments have similar distributions across the genome in ATAC-seq experiments [**Supplementary Table 3**]. Such evidence indicated that the majority of the captured fragment represents the open chromatin signal that can be derived from the Tn5 insertions in ATAC-seq experiments. To better represent the Tn5 insertion event, we shifted each end of the non-redundant uniquely mapped read pair +4 bp/−5 bp on the positive/negative strands to define the Tn5 insertion position and then further extended 75 bp in both directions around the Tn5 insertion position. By applying this strategy, one non-redundant uniquely mapped read pair was divided into two SE fragments [PE(asSE)], and the sequencing depth doubled compared to that of traditional analysis methods that manage PE fragments without shifting [PE(noShift)] [**Figure 3A**].

**Figure 3.**
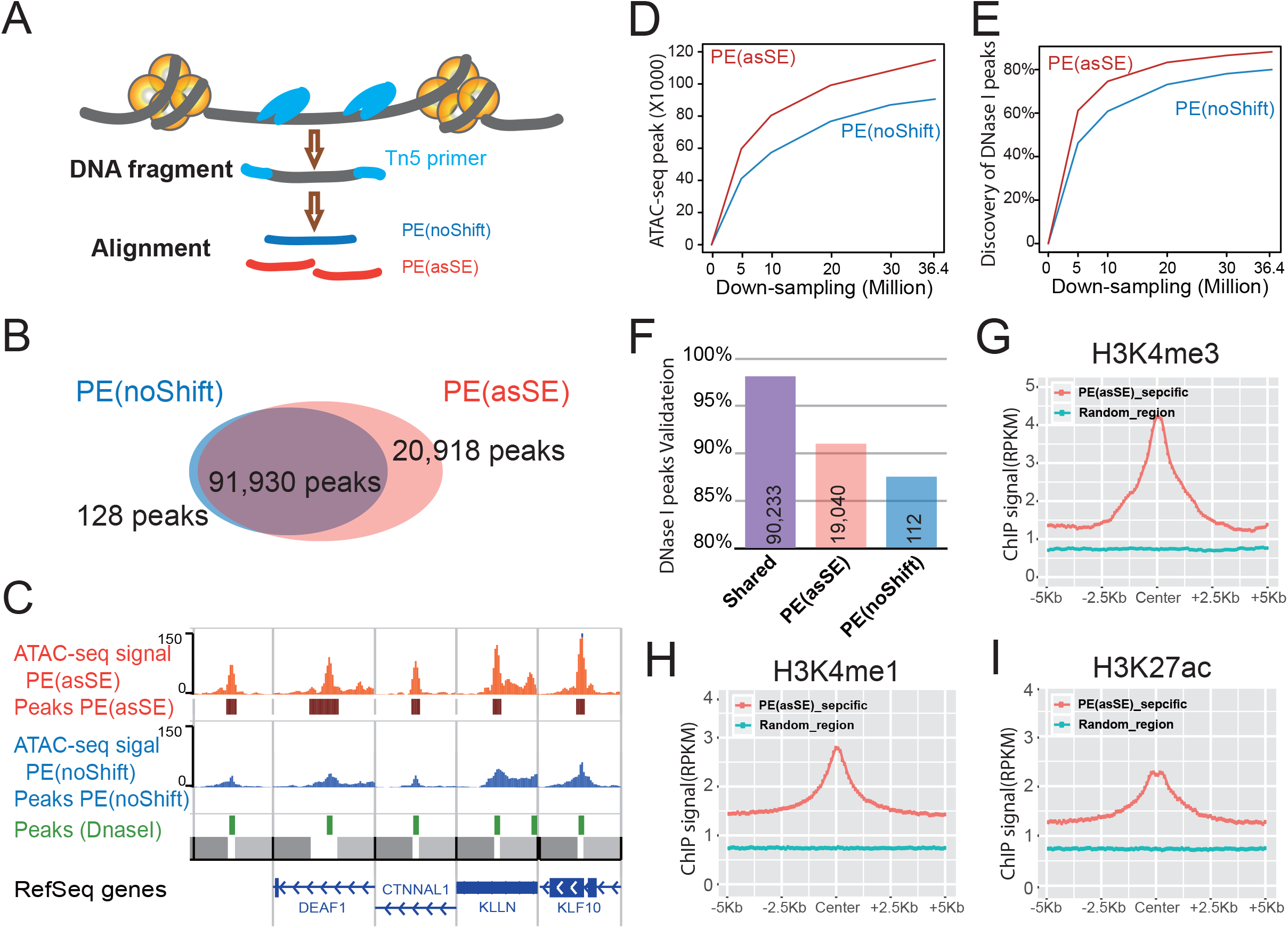
Comparison of the peak-calling strategy of AIAP [PE(asSE)] and the classic method [PE(noShift)]. **A).** Schematic representation of PE(noShift) and PE(asSE). PE(noShift) representing the complete DNA fragment insertion was generated as in previous studies. PE(asSE) representing one DNA fragment insertion was generated by AIAP. **B)**. Overlap of identified open chromatin regions (peaks) between PE(noShift) and PE(asSE) in the GM12878 cell line. **C)**. GM12878 ATAC-seq data were independently processed as PE(asSE) (red) and PE(noShift) (blue) and visualized with the WashU Epigenome Browser with DNase-seq peaks of GM12878 (green). Five regions were randomly selected to contain the peaks identified only in PE(asSE) mode but not in PE(noShift) mode. **D)**. Number of ATAC-seq peaks identified at different sequencing depths by random sampling of the GM12878 Omni-ATAC-seq data. **E)**. Discovery rate of GM12878 DNase-seq peaks by PE(asSE) and PE(noShift) at different sequencing depths (same random sampling as in **D**). **F)**. Percentage of shared, PE(asSE)-specific, and PE(noShift)-specific OCRs validated by merged DNase-seq data from ENCODE. **G, H, I)**. PE(noShift)-specific OCRs were enriched significantly with active histone modifications H3K4me3 (**G**), H3K4me1 (**H**), and H3K27ac (**I**) when compared to 20,000 regions randomly selected from the genome (green).

To validate the sensitivity of our analysis strategy, we downloaded published ATAC-seq data of GM12878 cells generated by the Greenleaf laboratory with the Omni-ATAC-seq protocol[31]. We first processed the data by following the classical method based on non-redundant uniquely mapped PE reads [PE(noShift)] and performed peak calling. In PE(noShift) mode, we identified 92,058 narrow peaks. In parallel, we applied AIAP to process the same data in PE(asSE) mode and performed peak calling with identical parameters (see Methods), and 112,848 peaks were identified. Compared to PE(noShift) mode, PE(asSE) mode reported ~99.9% of PE(noShift) peaks and identified ~23% additional peaks (20,918) [**Figure 3B**]. By visually inspecting the signal density on the genome browser, we noticed that most of the PE(asSE)-specific peaks overlapped with known ENCODE DHSs [**Figure 3C**]. We examined the ATAC-seq peaks and known GM12878 DHSs obtained from the ENCODE data portal. We noticed that PE(asSE) mode identified more OCRs at different sequencing depths [**Figure 3D**]. When analysing the full GM12878 Omni-ATAC-seq dataset, PE(noShift) mode identified ~80% of DHSs, and PE(asSE) mode identified ~85% of DHSs [**Figure 3E**]. We further used merged DHSs of 95 cell lines to measure the specificity of identified OCRs. We found that nearly 98% of common peaks identified by both PE(asSE) and PE(noShift) modes overlapped with known DHSs. A total of 19,040 of 20,918 peaks identified only by PE(asSE) mode overlapped with known DHSs, and 112 of 128 peaks identified only by PE(noShift) mode overlapped with known DHSs [**Figure 3F**]. These results suggest that PE(asSE) mode can greatly improve the sensitivity of the OCR discovery.

OCRs are generally considered regulatory elements that are enriched for specific histone modifications. We downloaded ChIP-seq data of GM12878 histone modifications (H3K4me3, H3K4me1, and H3K27ac) that are associated with promoter and enhancer activities to validate the functionality of PE(asSE)-specific OCRs. Compared to randomly selected genomic regions, the PE(asSE)-specific OCRs were highly enriched in all active histone modifications [**Figure 3G-I**]. These results suggest that the PE(asSE)-specific OCRs are functional regulatory elements rather than false positives. We further utilized AIAP to analyse the ATAC-seq data of multiple tissues, and compared to the classic PE(noShift) mode, the PE(asSE) mode resulted in a 28% to 55% increase in sensitivity when processing the ATAC-seq data [**Supplementary Table 4**]. These results indicated that AIAP can dramatically enhance the sensitivity of OCR discovery with high specificity.

### AIAP improved the sensitivity of DAR identification

Chromatin accessibility is dynamically associated with cellular responses to developmental cues, disease progression, and environmental stimuli. The identification of DARs has become an important approach to monitor the activity changes of regulatory elements. Since AIAP dramatically increased the sensitivity of OCR discovery, we further tested the sensitivity of AIAP in identifying DARs. We downloaded ATAC-seq dataset of mouse liver embryos at 11.5 days (E11.5d) and at the postnatal 0-day (P0d) stage from the ENCODE data portal and processed these data in both PE(asSE) and PE(noShift) modes. As expected, PE(asSE) mode identified 30% more OCRs than PE(noShift) mode [**Supplementary Table 4**]. To test the sensitivity of DAR identification, we used the complete set of OCRs identified in PE(asSE) mode and calculated the read counts based on both PE(asSE) and PE(noShift) modes [see Methods]. A total of 11,040 E11.5d-specific and 9,584 P0d-specific DARs were identified by both modes (shared DARs). We also identified 4,213 E11.5d-specific and 2,819 P0d-specific DARs by using only PE(asSE) mode [**Figure 4A**]. Correspondingly, only 107 E11.5d-specific and 72 P0d-specific DARs were found by using PE(noShift) mode. Compared to PE(noShift) mode, PE(asSE) mode resulted in an ~35% increase in DARs. We further tested AIAP on other tissues at two developmental stages and found that AIAP identified 32% to 168% more DARs in different tissues [**Supplementary Table 5**].

**Figure 4.**
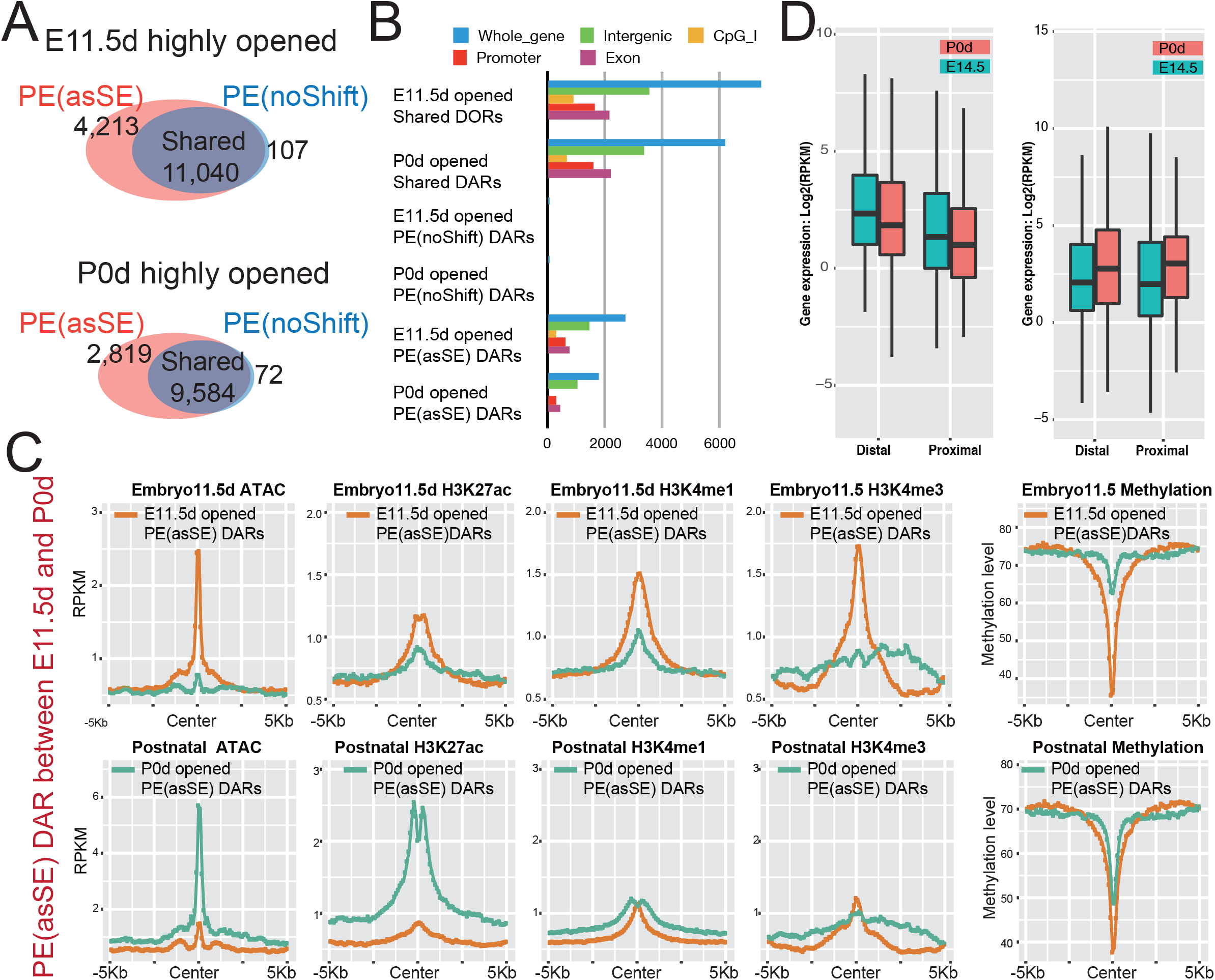
Comparison of the differential accessibility analysis of AIAP [PE(asSE)] and classic method [PE(noShift)]. **A).** Overlap of identified DARs between PE(noShift) and PE(asSE) when comparing embryonic 11.5-day (E11.5d) mouse liver to postnatal 0-day (P0d) mouse liver. **B)**. Genomic distribution of 21,624 shared, 7,032 PE(asSE)-specific, and 179 PE(noShift)-specific DARs. **C)**. Enriched epigenetic modifications (Left to right: ATAC-seq, H3K27ac ChIP-seq, H3K4me1 ChIP-seq, H3K4me3 ChIP-seq, and DNA methylation) on the PE(asSE)-specific DARs in the embryonic 11.5-day (Top) and postnatal 0-day stages (Bottom). **D)**. The expression of genes associated with PE(asSE)-specific E11.5d DARs (**Left**) and P0d DARs (**Right**) in proximal (2 kb around gene TSSs) and distal (2-20 kb around gene TSSs) in the two developmental stages.

We examined the genomic distribution of DARs and noticed that the distribution of PE(asSE)-specific DARs had a similar distribution as shared DARs: most DARs were located in intergenic and intronic regions, consistent with their potential enhancer functionality [**Figure 4B**]. Because dynamic changes in chromatin accessibility accompany the alteration in epigenetic modification synchronously[1, 8, 28, 33], we further used epigenetic data from the same samples to validate the accuracy of the DARs. We first checked the epigenetic modification around PE(asSE)-specific DARs. Compared to P0d-specific PE(asSE)-specific DARs, the E11.5d-specific DARs recruited highly active histone modifications associated with regulatory elements specifically at the 11.5-day stage of the embryo but not the postnatal 0-day stage, including H3K27ac, H3K4me1, and H3K4me3 [**Figure 4C**, **top**]. In contrast, the P0d-specific PE(asSE)-specific DARs recruited highly active histone modification H3K27ac specifically at the postnatal 0-day stage but not H3K4me1 or H3K4me3 modification [**Figure 4C**, **bottom**]. We also noticed that the E11.5d-specific DARs were significantly less methylated at the 11.5-day stage of the embryo, but P0d-specific DARs remained methylated at the E11.5d stage. However, E11.5d-specific DARs were still un-methylated at the later postnatal stage. The observation of the loss of DNA methylation on regulatory elements during embryo development is consistent with that of a previous study[34]. The epigenetic modifications of shared DARs showed very similar patterns to the PE(asSE)-specific DARs [Supplementary **Figure 3**]. We further examined the expression of genes around identified DARs. The DARs were assigned to the nearest gene based on distance and were classified into a proximal group (2 kb around the TSS) and a distal group (2-20 kb around the TSS). We noticed that the expression of genes around PE(asSE) embryo DARs was downregulated during liver development. In contrast, the expression of genes around postnatal DARs was upregulated at the same time [**Figure 4D**].

## DISCUSSION

AIAP is a new tool to perform quality assurance and downstream analysis of ATAC-seq data. AIAP provides a complete data processing solution for ATAC-seq data and provides a web-based QC report. Using public datasets, we systematically tested the QC metrics of ATAC-seq data and established key QC metrics of ATAC-seq data. We determined that the reads under peak ratio (RUPr), promoter enrichment (ProEn), and background noise (BG) were important measurements to estimate the quality of ATAC-seq data. We found that the RUPr and ProEn reflect the ATAC-seq signal enrichment, and BG indicates the overall background noise of the data. By combining these QC metrics, we obtained an accurate estimation of the quality of ATAC-seq data. We used AIAP to process 54 mouse ATAC-seq datasets to test and evaluate the QC metrics and generate the range of the QC metrics [**Supplementary Table 2**]. These ranges of QC metrics can be used as a reference to evaluate the success of ATAC-seq experiments.

We optimized the widely used classic analysis methodology and specifically used pair-end as single-end (PE(asSE)) mode to process the PE sequenced ATAC-seq data. AIAP aligns the PE ATAC-seq data in PE mode to increase the alignment accuracy, and the BAM file is further processed in SE mode for downstream analysis. In PE(asSE) mode, AIAP doubles the sequencing depth and dramatically increases the sensitivity of OCR identification. In our test, AIAP identified 20-40% more OCRs than the widely used classic analysis method. We further used corresponding DHS data and histone modification data to validate the specificity of newly identified OCRs by AIAP; the majority of the novel OCRs identified by AIAP were independently validated with DHSs and enriched for active histone modifications. Such a result indicates the high true-positive rate resulting from the AIAP analysis strategy.

We also suggest that the PE(asSE) strategy can improve not only the sensitivity of discovering OCRs but also the sensitivity of discovering identical DARs. By using ENCODE ATAC-seq data of liver embryo development, we found that AIAP can identify 32% to 168% more chromatin DARs than the classic PE(noShift) mode. We further indicated that the novel DARs identified by AIAP were enriched in active histone modifications at different developmental stages. The embryonic 14.5 day-specific DARs were lowly methylated, and the H3K27ac signals were significantly enriched only in the 14.5-day embryo stage but not in the postnatal 0-day stage. In contrast, the postnatal 0-day-specific DARs were highly methylated in 14.5-day embryo stage without active histone modifications and became minimally methylated and recruited strong active H3K27ac signals in the postnatal 0-day stage. We also noticed that the expression of genes around the developmental stage-specific DARs was associated with the openness of DARs, as other studies reported[35, 36]. These results suggested that AIAP can greatly improve the sensitivity to identify DARs with high specificity.

Finally, we compiled AIAP into a Docker/Singularity image to facilitate the easy operation of AIAP on high-performance computing clusters. AIAP can complete QC checking and file processing in 2-3 hours for one typical ATAC-seq dataset. AIAP supports multiple genome assemblies, including human (hg19 and hg38), mouse (mm9 and mm10), and rat (rn6). Additionally, each step for QC and data processing is componentized and can be called by advanced users to build pipelines for specialized applications, and different genome assemblies can be easily and directly compiled for ATAC-seq data processing in other species.

## AVAILABILITY

The software, source code, and documentation are freely available at https://github.com/Zhang-lab/ATAC-seq_QC_analysis.

## SUPPLEMENTARY DATA

Supplementary Table 1, 2, 3, 4, 5

Supplementary Figure 1, 2, 3

## FUNDING

This work was supported by the National Institutes of Health [U24ES026699, U01HG009391, and R25DA027995]; and the Goldman Sachs Philanthropy Fund [Emerson Collective]. Funding for open access charge: National Institutes of Health.

## CONFLICT OF INTEREST

All authors declare no conflicts of interest.

## Notes

https://github.com/Zhang-lab/ATAC-seq_QC_analysis

