## Supplementary material for "Improving ATAC-seq Data Analysis with AIAP, a Quality Control and Integrative Analysis Package": S-Table-figure

* To whom correspondence should be addressed.

**Supplementary Table**

**Supplementary Table 1**: List of data used in manuscript

**Supplementary Table 2**: QC metrics of 70 mouseENCODE ATAC-seq data

**Supplementary Table 3**: DNA Insertion fragments distribution under the peak

|  | **Forebrain** | **Intestine** | **Kidney** | **Liver** | **Lung** | **Stomach** |
| --- | --- | --- | --- | --- | --- | --- |
| Sample ID |  |  |  |  |  |  |
| Total # of fragments | 24914597 | 22141337 | 26597998 | 16120650 | 33646778 | 17254895 |
| Short fragments(< 38bp) | 25334 | 36920 | 106029 | 21373 | 55781 | 31901 |
| Medium fragments (38-150bp) | 8647206 | 9166167 | 13669940 | 5924854 | 12878709 | 8043302 |
| Long fragments (>150bp) | 16242057 | 12938250 | 12822029 | 10174423 | 20712288 | 9179692 |
| RUP^1^ (Medium frags) | 1860344 | 898813 | 1488011 | 1412188 | 3230218 | 534505 |
| RUP^1^ (long frags) | 2804650 | 1235315 | 1174004 | 2060910 | 5062100 | 456521 |
| RUPr^2^ (Medium frags) | 21.51% | 9.81% | 10.89% | 23.83% | 25.08% | 6.65% |
| RUPr^2^ (long frags) | 17.27% | 9.55% | 9.16% | 20.26% | 24.44% | 4.97% |

1) RUP: Reads Under Peaks; 2) RUPr: Reads Under Peaks ratio

**Supplementary Table 4**: Peak-calling comparison between PE(asSE) and PE(noShift)

|  | **Forebrain** | **Intestine** | **Kidney** | **Liver** | **Lung** | **Stomach** |
| --- | --- | --- | --- | --- | --- | --- |
| Sample ID | ENCLB042MOW | ENCLB362STB | ENCLB497HBT | ENCLB303HH | ENCLB080OEI | ENCLB490MG |
| OCRs # in PE(noShift) | 39127 | 22644 | 25437 | 23044 | 58283 | 17012 |
| OCRs # in PE(asSE) | 49993 | 33548 | 39259 | 28851 | 77796 | 23882 |
| Shared OCRs | 38830 | 22474 | 25209 | 22924 | 57987 | 16881 |
| PE(asSE)-specific OCRs | 11163 | 11074 | 14050 | 5927 | 19809 | 7001 |
| Increase | 28.75% | 49.27% | 55.73% | 25.85% | 34.16% | 41.47% |

**Supplementary Table 5**: Differential analysis comparison between PE(asSE) and PE(noShift)

|  | **Forebrain** | **Intestine** | **Kidney** | **Liver** | **Lung** | **Stomach** |
| --- | --- | --- | --- | --- | --- | --- |
| Sample ID-1 | ENCLB252ZLC | ENCLB069VWJ | ENCLB087XNG | ENCLB200ODB | ENCLB517ADM | ENCLB880WNY |
| Sample ID-2 | ENCLB312MJN | ENCLB325XDP | ENCLB299QNB | ENCLB555EYH | ENCLB224FWZ | ENCLB322JPD |
| Sample ID-3 | ENCLB042MOW | ENCLB362STB | ENCLB678YRF | ENCLB303HHQ | ENCLB080OEI | ENCLB490MGZ |
| Sample ID-4 | ENCLB558DNK | ENCLB199HRA | ENCLB497HBT | ENCLB282VPH | ENCLB071NVJ | ENCLB105JML |
| Total # of OCR | 126784 | 77483 | 83371 | 89128 | 129812 | 97233 |
| DARs # in PE(noShift) | 5113 | 7131 | 700 | 11121 | 11177 | 451 |
| DARs # in PE(asSE) | 8464 | 10808 | 1646 | 14746 | 16402 | 1211 |
| Shared DARs # | 5084 | 7009 | 691 | 11051 | 11059 | 448 |
| PE(asSE)-speific DAR | 3351 | 3677 | 946 | 3625 | 5225 | 760 |
| Increase | 65.54% | 51.56% | 135.14% | 32.60% | 46.75% | 168.51% |

**Supplementary Figures**

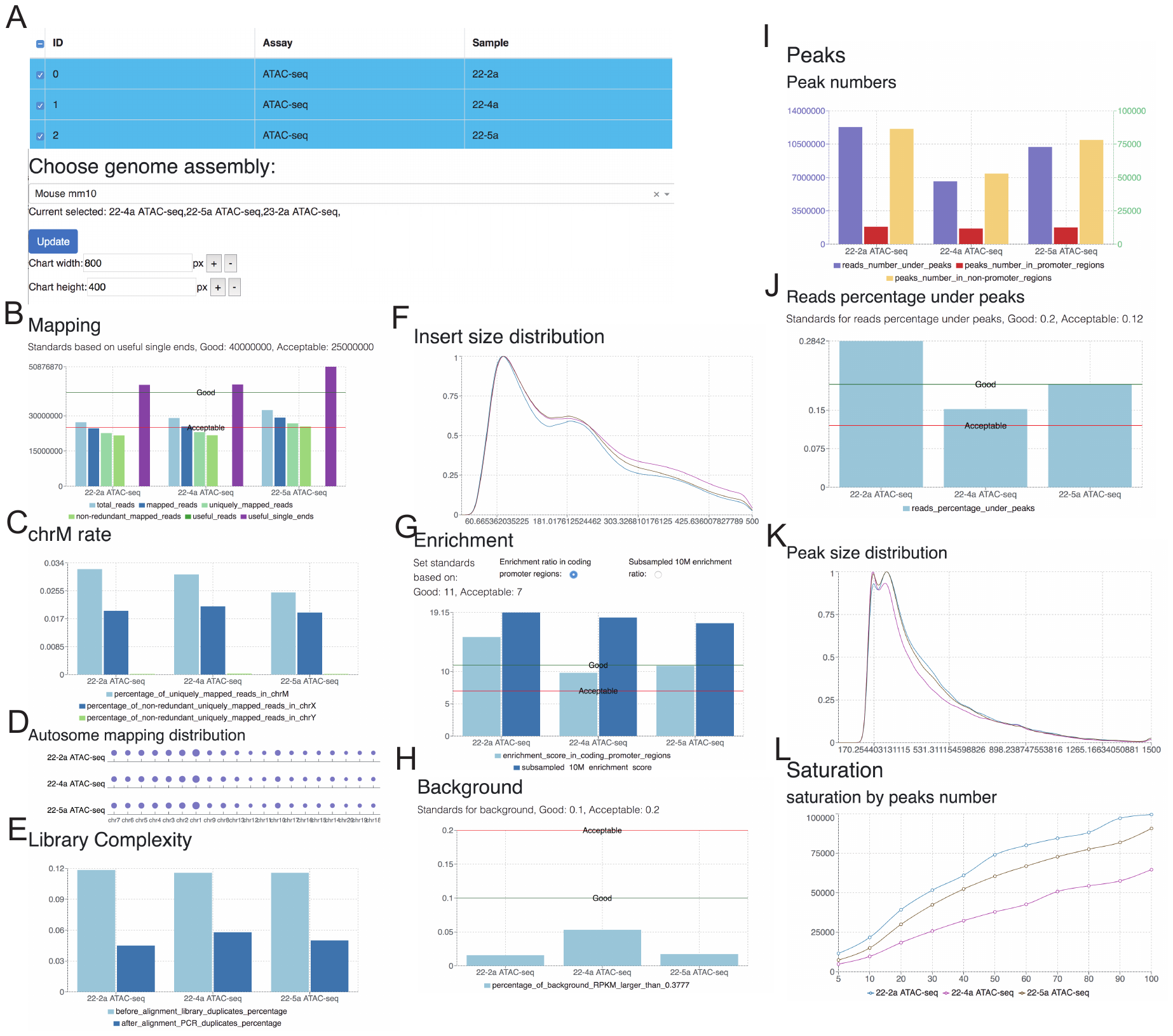

**Supplementary Figure 1. ATAC-seq QC report visualized by QAviewer for 3 selected ATAC-seq datasets. A).** Summary of data and figure configuration. **B)**. Mapping summary. **C)**. Percentage of uniquely mapped reads on chrM, chrX, and chrY. **D)**. Percentage of uniquely reads on all the autosome. **E)**. Library complexity estimation by FastQC (before alignment) and PreSeq (after alignment). **F)**. The length distribution of inserted DNA fragments. **G)**. Promoter Enrichment and Subsampling Enrichment. **H)**. Background. **I)**. The number of reads under the peak (Left), and the number of peaks in gene’s promoter (Red) and the number of peaks out of promoter (Yellow). **J)**. The Reads Under Peak ratio (RUPr). **K)**. The length distribution of all peaks. **L)**. Saturation analysis result.

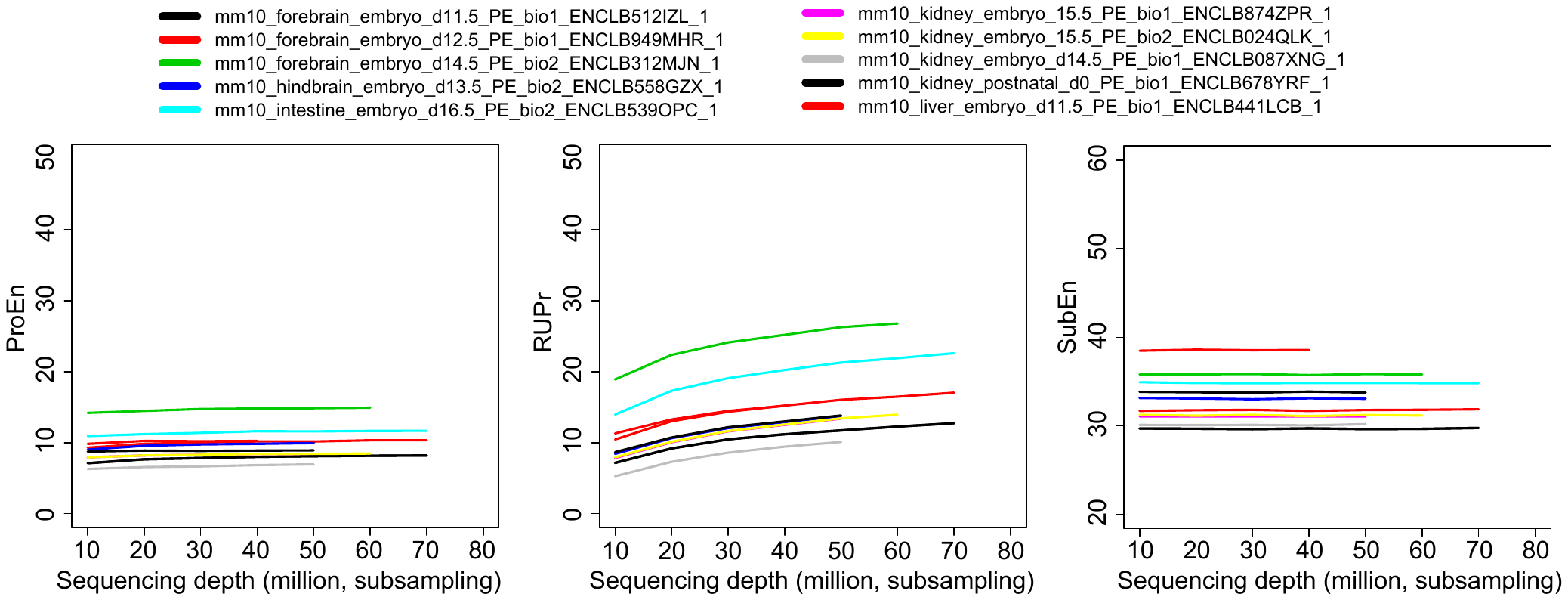

**Supplementary Figure 2. Key QC metrics with subsampling test. The x-axis is the subsampling useful single ends. Left).** Promoter Enrichment. **B)**. Read Under Peak ratio. **C)**. Subsampling Enrichment

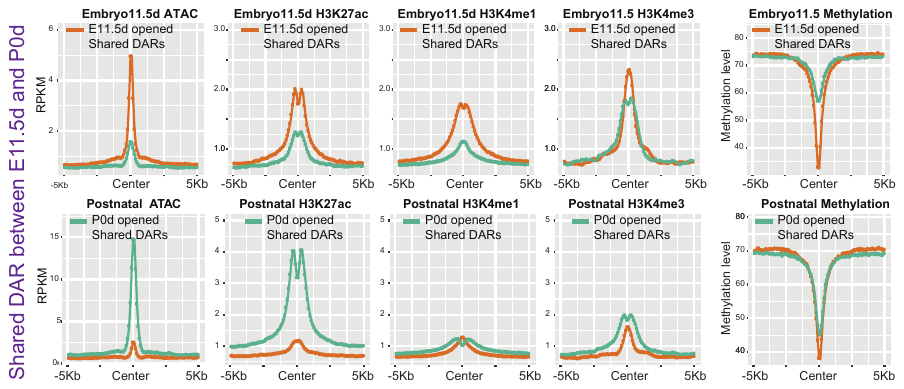

**Supplementary Figure 2.** Enriched epigenetic modifications (left to right: ATAC-seq, H3K27ac ChIP-seq, H3K4me1 ChIP-seq, H3K4me3 ChIP-seq, DNA methylation) on the shared DARs in embryo 11.5 day (Top) and postnatal 0 day (Bottom).
